# Misaligned plastic and evolutionary responses of lifespan to novel carbohydrate diets

**DOI:** 10.1101/2022.11.14.516515

**Authors:** Vikram P. Narayan, Nidarshani Wasana, Alastair J. Wilson, Stephen F. Chenoweth

## Abstract

Diet elicits varied effects on longevity across a wide range of animal species. For example, diets low in protein and high in carbohydrate typically extend lifespan while diets high in protein tend to reduce it. Although studies have also shown that diet-induced lifespan changes can persist through transgenerational plasticity, whether such changes lead to evolutionary shifts in lifespan remains unclear. In this study we combine experimental evolution and phenotypic plasticity assays to address this gap. Using *Drosophila serrata*, we investigated the evolutionary potential of lifespan in response to four novel diets spanning a carbohydrate-protein gradient. We also examined developmental plasticity effects using a set of control populations that were raised on the four novel environments. Our results show that although lifespan evolved in response to changes in dietary carbohydrate concentration, the plastic responses for lifespan differed from the evolved responses. The direction of the evolved response (increased lifespan) observed on low carbohydrate diets was in the opposite direction to the plastic response (decreased lifespan). Our results imply that plastic responses to low carbohydrates can be maladaptive for lifespan and misaligned with the evolved responses, laying the groundwork for future investigations of carbohydrate contributions to evolved and plastic effects on lifespan.

## Introduction

Diet is a fundamental component of an organism’s environment, responsible for physiological and morphological adaptations [1]. Variation in dietary macronutrient balance, specifically the ratio of protein to carbohydrate, has been shown to have strong influences on lifespan. For example, in insects, diets low in protein and high in carbohydrate typically extend lifespan while diets high in protein tend to reduce it [2–8]. Further, lifespan is maximised at different macronutrient ratios in males and females indicating fundamental differences in resource acquistion and allocation between the sexes [9–17]. To date, most of what is known about the effects of diet on longevity, especially its extension, comes from studies manipulating nutrient content within a single generation [18]. A smaller number of studies have shown that diet-induced lifespan changes can persist through transgenerational plasticity for a small number of generations [19–24]. Experimental evolution has also revealed that lifespan can evolve during adaptation to novel diets [23, 24]. However, it is not yet known whether lifespan changes seen during experimental evolution mirror those observed in studies of phenotypic plasticity.

Broadly, it is well recognised that genotypic evolution and phenotypic plasticity are two key mechanisms by which populations respond to environmental change, however, their relative contributions remain unresolved [25–28]. For instance, phenotypic plasticity is generally assumed to be adaptive, allowing the phenotype to move in the direction of higher fitness. However, plasticity can also be maladaptive, moving the phenotype in the opposite direction when organisms are exposed to unusual or novel environments [29, 30]. In the context of diet and lifespan, recent studies in *Drosophila* have demonstrated the potential for lifespan to evolve genetically under different dietary conditions, but the degree of alignment of evolutionary with plastic responses to diet remains unclear [23, 24]. In this study we combine experimental evolution and phenotypic plasticity assays to bridge this gap in our understanding. Long-term adult health and lifespan is shaped by both current diet and past dietary history, where persistent long-term effects of nutrition transmitted from parent to offspring are known to program lifespan [31]. These effects, including caloric or dietary restriction (DR), can range from being instantaneous to inter-generational in nature [32]. In *Drosophila*, flies switched from DR to an *ad libitum* diet later in life immediately adopted the survival pattern of the group kept on the *ad libitum* diet treatment [33]. In murine models, switches between DR and unrestricted feeding have shown mixed results but the results of a meta-analysis do suggest the presence of a lasting ‘memory’ of diet [34].

Disproportionate dietary sucrose has been used to create a fly model of an obesogenic and unhealthy diet that bares relevance to modern human diets, however different hypotheses for the evolution of lifespan under a high-carbohydrate diet have rarely been tested, and a capacity for lifespan to evolve in response to macronutrients is yet to be established for many model systems (but see [23, 24]). Diet concentrations have been shown exhibit a bell-shaped relationship with lifespan, ranging from under- to over-nutrition (Figure 1). Much of our current understanding of how diet alters lifespan comes from studies using model organisms where diet has been manipulated by changing the proportions of protein relative to carbohydrate. However, little is known about the effects of DR and carbohydrate diets on longevity phenotypes in response to changing carbohydrate concentrations while keeping protein concentrations constant.

**Figure 1.**
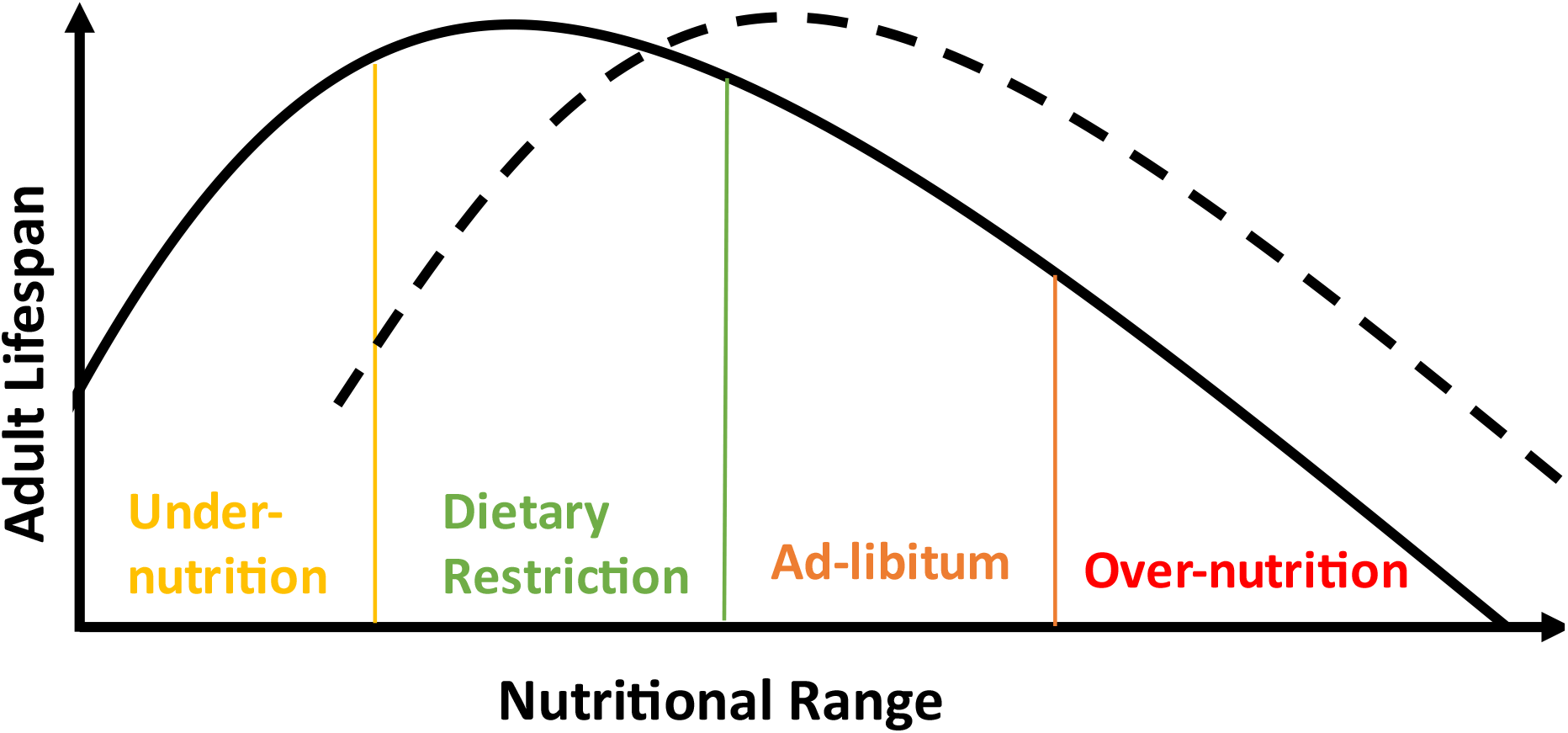
Lifespan reaction norm to diet. Diet nutritional content has a convex shaped relationship with lifespan, ranging from under-nutrition, dietary restriction and maximal lifespan to ad-libitum feeding and over-nutrition. The lifespan reaction norm can shift or change shape (dashed line) in response to genetic or environmental changes. For instance, dietary restriction effects can lead to increased lifespan on the focal curve but undernutrition on the dashed curve. Adapted from [35].

In this study, we used an outbred population of *Drosophila serrata* [36] that has adapted for over 200 generations to laboratory culture in a diet where the carbohydrate content is 1.5 times that of the protein content. To determine if the response of lifespan to changing carbohydrates concentrations follow the same patterns as changing protein concentrations, we created four novel diets where we varied the carbohydrate content by manipulating sugar while keeping protein (yeast) content constant. For 30 generations we exposed replicates of the founding population to these four novel diets spanning a carbohydrate-protein gradient to detect evolved responses in lifespan. To elucidate developmental plasticity effects, we concurrently assayed lifespan in a set of control populations that were exposed to the four novel environments. This approach allows us to assess the extent to which plastic responses of lifespan mirrored adaptive responses observed following experimental evolution. We expected to find the evolution of extended lifespan on low carbohydrate diets and reduced lifespan on the high carbohydrate diets for both sexes, and the plastic responses to resemble the evolved responses. While we found both high- and low-carbohydrate diets to elicit plastic and evolved effects on lifespan, the plastic responses for lifespan to developmental diets bore little resemblance to the evolved responses on evolutionary diets.

## 2. Methods

### 2.1 Fly Husbandry

Flies used to set up the experimental evolution population experiment were all derived from a wild-type, outbred and long-term laboratory-adapted population that was founded from 102 inbred lines of the *D. serrata* Genomic Reference Panel established from a single natural population in Brisbane, Australia [36]. When the experiment began, the population had been maintained at The University of Queensland for 150 generations at a large population size, on a 12-h:12-h light: dark cycle at 25 °C on standard 1.5 carbohydrate (sugar): protein (yeast) (SY) diet.

*D. serrata* flies from the outbred and long-term laboratory-adapted population were evolved on five distinct SY diets (0.5, 1.0, 1.5, 2.0 and 2.5 SY) for 30 generations before the start of the experiment (specific diet characteristics are given in supplementary Table S1). There were four replicate populations per diet treatment and each replicate consisted of four mixed-sex population bottles that was established from 60 adult flies each generation.

### 2.2 Experimental design

Assay flies were provided with one of the five experimental evolution diets (Figure 2). Two generations of common garden were used to remove parental effects from the SY diet treatments before lifespan and survival of the third-generation offspring was measured. The solid blue arrow shows the transfer from the ancestral diet to every other developmental diet and differences in lifespan here provide evidence of evolutionary changes to lifespan across generations. The dotted blue arrow shows the transfer of every evolutionary diet back to the ancestral diet and differences in lifespan among diet treatments here demonstrate developmental plasticity effects of lifespan.

**Figure 2.**
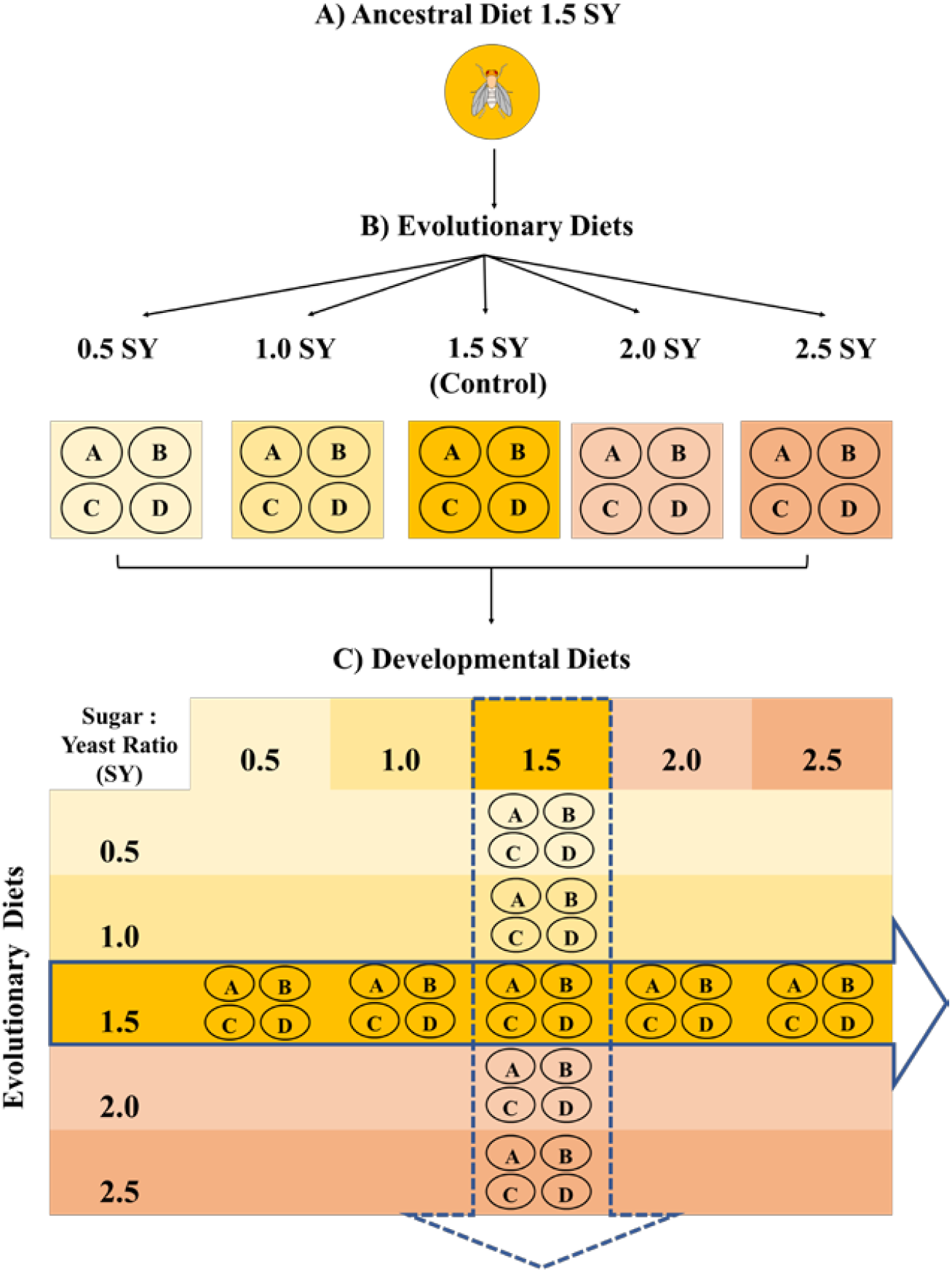
Experimental Evolution Design where A) *D. serrata* flies from an outbred and long-term laboratory-adapted population were evolved on five distinct diets with different sugar content as adults for 30 generations (supplementary table S1). B) Flies from each diet in the experimental evolution experiment were kept as four independent replicates. C) To remove any parental effects from the evolution diet treatments, experimental flies were subjected to two generations of common garden before we used ten replicate vials with ten flies per replicate for each evolution diet (n = 400) to test for evolved and plastic lifespan responses in third-generation offspring. The solid blue arrow shows the transfer of the ancestral diet replicate populations lines to every other developmental diet and the dotted blue arrow shows the transfer of replicates from every evolutionary diet back to the ancestral diet.

We used ten replicate vials with ten flies each per bottle for each evolution diet × assay diet combination (N = 400). Sample sizes are given in supplementary tables S2 and S3. Flies were maintained at 25°C and 60% humidity, on a 12:12 h light: dark cycle. Survival was checked every 3-4 days and flies were tipped into fresh food vials until all flies had died. On these occasions, dead flies were counted and removed from vials.

### 2.3 Statistical Analysis

PROC GLIMMIX (SAS software v9.4) was used to analyse treatment effects on adult lifespan in *D. serrata* according to the split-split-plot experimental design outlined in [37]. We used a mixed effects linear model to analyse the evolved and plastic effects of diet on lifespan. Models were fit a categorical ‘diet’ effect that included the five diets (0.5, 1.0, 1.5, 2.0, and 2.5 SY). Given the sex differences in *D. serrata* lifespan, different models were fit to examine the effects of evolutionary and developmental diets with sex as a fixed effect to control for and detect any sex-specific effects.

The model for plastic responses contained ‘*developmental diet*’ as the fixed effect:

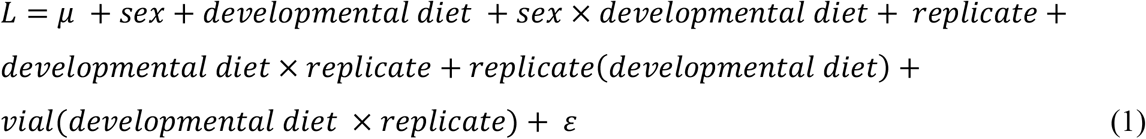

where lifespan *L* is measured in days, *μ* is mean lifespan, *developmental diet* is the fixed main effect of diet, *replicate* is a random effect describing variation among the four replicate (control, 1.5 SY diet) populations from the evolution experiment, and *ε* is the residual error.

The model for evolved responses contained ‘*evolutionary diet*’ as a fixed effect:

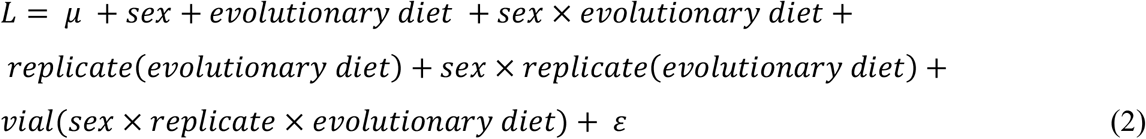

where lifespan *L* is measured in days, *μ* is mean lifespan, *evolutionary diet* is the fixed main effect of diet, *replicate*(*evolutionary diet*) is a random effect of replicate population nested within diet, and *ε* is the residual error.

For both models, we used a Satterthwaite approximation to calculate the denominator degrees of freedom via the “ddfm=SAT” option in SAS software when testing significance of the diet effect. We present effect sizes as least square means and used Tukey’s HSD to test for differences between specific diets (α = 0.05).

## 3. Results

### 3.1 Plasticity of lifespan in response to diet

We found that relative to the intermediate 1.5 SY diet, both high- and low-carbohydrate diets elicited plastic effects on lifespan. Lifespan responded to changes in nutritional environments for both sexes resulting in a bell-shaped relationship between lifespan and increasing carbohydrate content (Figure 3). Lifespan responded to the novel diet treatments, with significant main effects of *sex* (*F*_1,371.6_ = 476.9, *p* = 1.32e^−68^) and *developmental diet* (*F*_4,12.1_ = 6.54, *p* = 4.90e^−03^) detected, but not *sex* × *developmental diet* (*F*_4,371.6_ = 1.82, *p* = 0.12) (Figure 3). The plastic effects resemble a bell-shaped relationship between lifespan and increasing carbohydrate content with lifespan in both males and females maximised around the intermediate 1.5 SY developmental diet (Figure 3). Instead of increased lifespan on low SY diets as expected, there was a nonlinear relationship between developmental diets and lifespan that negatively affected adult survival (Figure 3, S2). By comparison, for males, higher lifespan was observed over a broader range of low SY developmental diets, albeit with a tendency for lifespan to still peak on intermediate 1.5 SY developmental diets (Figure 3).

**Figure 3.**
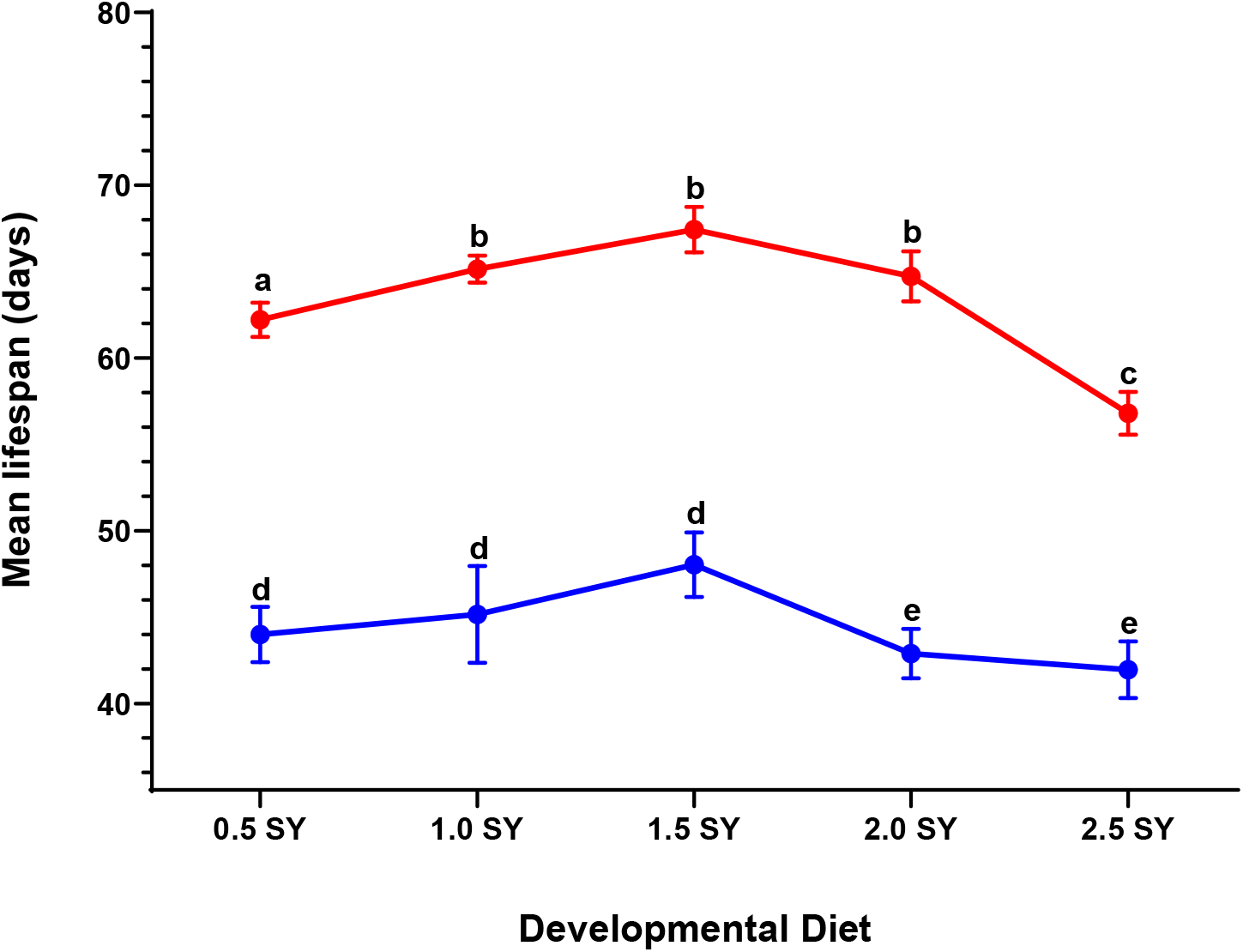
*D. serrata* mean lifespan on developmental diets. Mean adult female (red bars) and male (blue bars) life span measured in days is shown for the 1.5 SY evolutionary diet transferred to every other developmental diet. Error bars represent the 1 S.E. of the mean. Letters indicate significance in planned comparisons between 1.5 SY control and novel diets within sexes.

Both high and low carbohydrate developmental diets significantly decreased longevity in female *D. serrata*. In females, post-hoc comparisons relative to the 1.5 SY ancestral diet showed that, flies raised on 0.5 and 2.5 SY developmental diets lived significantly shorter lives (0.5 SY vs. 1.5 SY: t = −2.42, *p* = 0.02; 2.5 SY vs. 1.5 SY: t = 4.90, *p* = 1.40e^−05^; Figures 3; S2A and Table S6). Mean lifespan was not significantly different on the 1.0 and 2.0 SY developmental diets which were closer to the control 1.5 SY diet (1.0 SY vs. 1.5 SY: t = −1.06, *p* = 0.30; 1.5 SY vs. 2.0 SY: t = 1.26, *p* = 0.23; Figures 3; S2A and Table S6). The significant plastic effects of developmental diets for lifespan could be driven by flies on both lower and higher SY diets living less and ageing faster than flies from populations closer to the ancestral 1.5 SY diet (Figures 3 and S2A).

Post-hoc comparisons for males showed that mean lifespan was not significantly different on any of the low SY developmental diets relative to the ancestral 1.5 SY control diet (0.5 SY vs. 1.5 SY: t = −1.90, *p* = 0.07; 2.5 SY vs. 1.5 SY: t = 0.95, *p* = 0.35; Figures 3, S2B, Table S3 and S7). However, compared to the 1.5 SY ancestral diet, male flies raised on 2.0 and 2.5 SY developmental diets lived significantly shorter lives (2.0 SY vs. 1.5 SY: t = 2.41, *p* = 0.02; 2.5 SY vs. 1.5 SY: t = 2.85, *p* = 0.01; Figure 3, S2B, Table S3).

### 3.2 Evolution of lifespan in response to diet

Having detected lifespan plasticity in response to the developmental diets, we then tested whether those developmental plastic responses matched evolved responses to the same evolutionary diets. After approximately 30 generations of experimental evolution, we detected significant main effects of *evolutionary diet* (*F*_4,15.0_ = 10.2, *p* = 3.00e^−04^) and *sex* (*F*_1,367.2_ = 544.8, *p* = 1.57e^−74^), but not *sex* × *evolutionary diet* (*F*_4,367.2_ = 1.47, *p* = 0.21). With an overall negative relationship between lifespan and carbohydrate content, we found that both high- and low-carbohydrate diets elicited evolutionary effects on lifespan (Figure 4). However, evolved responses to evolutionary diets did not resemble the pattern of plastic responses for lifespan on SY developmental diets.

**Figure 4.**
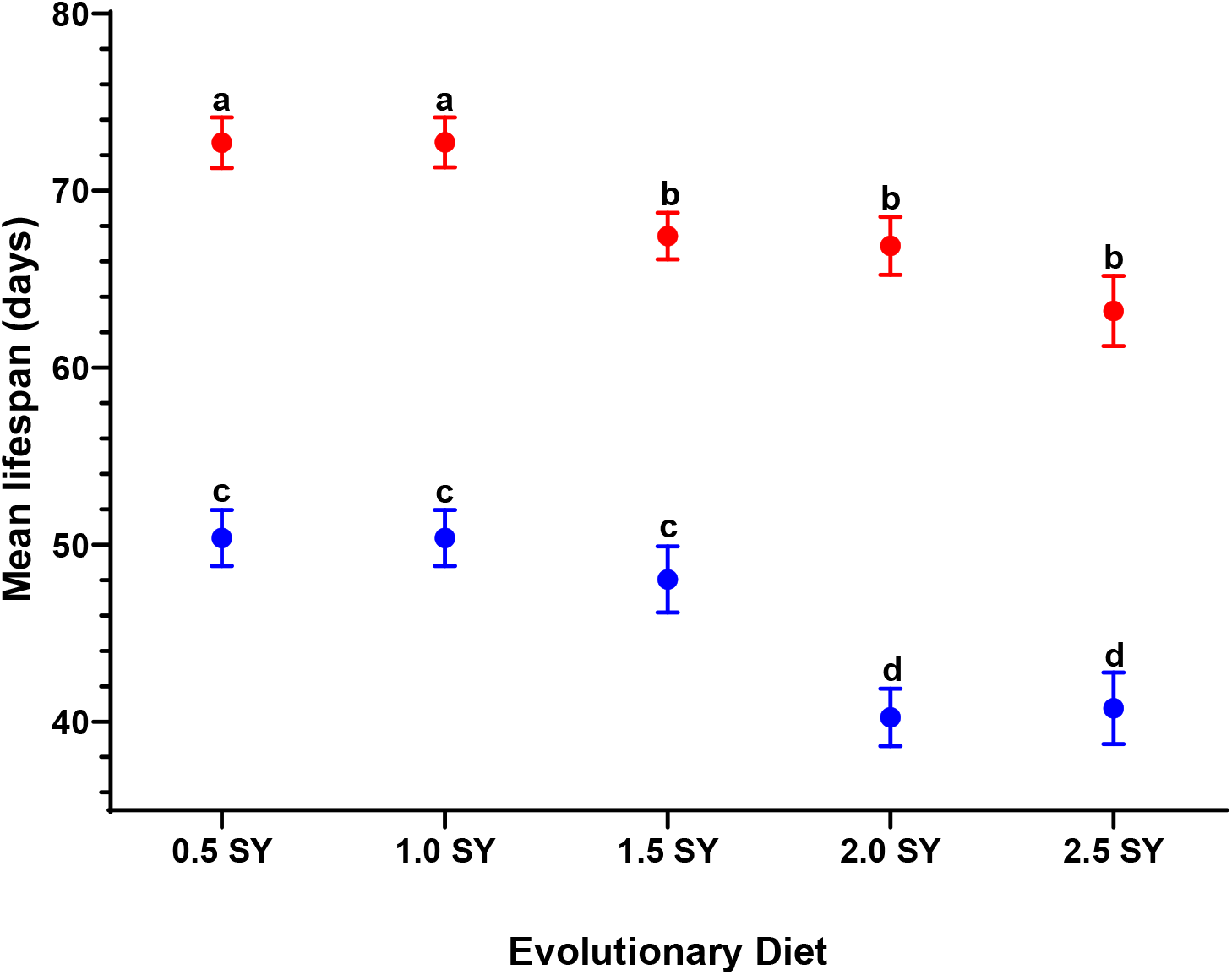
*D. serrata* mean lifespan on evolutionary diets. Mean adult female (red bars) and male (blue bars) life span measured in days is shown for every evolutionary diet transferred to the 1.5 SY ancestral diet. Error bars represent the 1 S.E. of the mean. Letters indicate significance in planned comparisons between 1.5 SY control and novel diets.

Post-hoc comparisons between females showed that flies evolved on lower SY diets of 0.5 and 1.0 SY lived on average 5.3, 5.8 and 9.5 days longer than flies evolved on 1.5, 2.0 and 2.5 SY diets respectively (Supplementary Table S2). Mean lifespan was significantly different on 0.5 and 1.0 SY diets (0.5 SY vs. 1.5 SY: t = 2.05, *p* = 0.047; 1.0 SY vs. 1.5 SY: t = 2.06, *p* = 0.046; Figures 3; S1A; Table S4). Mean lifespan was not significantly different on the 2.0 and 2.5 SY diet compared to the ancestral control diet of 1.5 SY (1.5 SY vs. 2.0 SY: t = 0.25, *p* = 0.80; 1.5 SY vs. 2.5 SY: t = 1.77, *p* = 0.09; Figures 4; S1A; Table S4). Female lifespan appeared more resilient to changes brought on by high SY evolution diets and evolution under high SY diets did not result in strong effects on female lifespan patterns.

In contrast to females, post-hoc comparisons showed that lifespan increases observed in male flies evolved on low sugar diets were not statistically significant (0.5 SY vs. 1.5 SY: t = 0.89, *p* = 0.38; 1.0 SY vs. 1.5 SY: t = 0.38, *p* = 0.38; Figures 4 and S1B). Males on the ancestral 1.5 SY diet lived on average 8 days longer than males on higher SY diets (1.5 SY vs. 2.0 SY: t = 3.19, *p* = 2.90e^−03^; 1.5 SY vs. 2.5 SY: t = 3.01, *p* = 4.60e^−03^; Figure 4; S1B; Table S2 and S5). Compared to females, males had less resilience to a high sugar diet leading to a reduction in lifespan. The significant effects of novel diet for lifespan could be driven by flies on lower SY diets living longer and ageing slower than flies from higher SY diet populations (Figures 3, S1A, Table S2, S4).

## 3. Discussion

### Dietary-based developmental plasticity affects lifespan

Our plasticity assay revealed a dietary optimum for male and female lifespan where developmental diet concentrations had a bell-shaped relationship with lifespan representative of the typical lifespan reaction norm to diet (Figure 1). We did not detect any lifespan extension under low SY developmental diets in females or males. Instead, compared to the ancestral diet, we found decreased lifespan on low SY developmental diets. In contrast to our findings, [24] demonstrated the typical expected DR effect, with a negative association between increasing dietary protein content and lifespan. However, DR responses even under protein manipulation are highly variable among different genotypes of *D. melanogaster* with some genotypes showing no or only little lifespan extension on a DR diet, with others showing decreased lifespan on a DR diet [38]. Furthermore, we manipulated sugar content while keeping yeast content the same, whereas most studies including [24] and [23] have manipulated yeast content. Our approach to diet manipulation was important for two reasons. First, carbohydrate-rich diets represent a fitting model of unhealthy, modern, human diets and the carbohydrate diets used in this experiment are a particularly relevant example of recent nutritional changes in many human populations with important implications for human health [39, 40]. Second, the effects of a high carbohydrate (sugar) manipulation in diet have not yet been studied extensively, though consistent with our findings, studies have shown that adults developed on high carbohydrate diets have reduced lifespan [41, 42].

Under life-history theory predictions of decreased fitness and lifespan, the plastic response to low SY diets in our study most likely corresponds to a “stress” that is expressed when populations have not yet adapted to this new dietary environment [43]. For example, gene expression plasticity evolved in *D. melanogaster* adapted to a medium enriched in salt and cadmium [43], but the direction of plasticity observed was in the opposite direction compared to the ancestral populations. Salt and cadmium represent novel and extreme environments seldom encountered by fruit flies outside laboratory settings. Such extreme environments suggest a likelihood of maladaptive plasticity and therefore the possibility of robust selection [44, 45], including on the plasticity of a trait [46]. Finally, although both sexes can demonstrate nutritionally plastic responses, in many species it is the larger sex (females in flies) that is more nutritionally plastic [47], a pattern consistent with the present study.

### Evolutionary adaptation to nutrient availability

Our experimental evolution experiments revealed that lifespan evolved in response to changes in dietary carbohydrate concentration and that the direction of the evolved response observed on low SY developmental diets was in the opposite direction to the plastic response. We found a strong positive effect of increased lifespan for evolution under low SY in females, whereas a prior experimental evolution study [23] found that females evolved under DR had lower lifespans compared to females that evolved on the standard diet. Another difference between this experimental evolution study and our study was that males evolved lower lifespans on high SY diets as opposed to no differences in male survivorship on any of the evolutionary diets in the previous study [24]. Flies evolved on higher carbohydrates, especially males, had shorter lifespans. According to previous studies, this suggests a preference or program in adults where even after the carbohydrate challenge has been removed, lifelong exposure to diets high in carbohydrates may affect many metabolic pathways such as those involved in glucose homeostasis, or cholesterol metabolism in offspring [48, 49].

Under specific circumstances, increased extrinsic mortality can select for the evolution of longer lifespans [50]. Dietary interventions that extend lifespan, such as DR effects induced by low SY diets can also be considered as a form of nutritional stress [51, 52], and may explain the potential to evolve longer lifespans. For example, exposure to heat stress in *C. elegans* requires strong anti-ageing mechanisms to survive, and this results in individuals with slower ageing [50]. Although constant dietary excess such as diets high in sugar have been known to accelerate ageing and reduce lifespan [18, 42, 53] it is still possible for such a diet to select for longer lifespans given a particular combination of conditions [54, 55]. Nutrient availability in response to distinct nutritional and metabolic demands of life-history strategies can cause organisms to experience alternative developmental programs [56]. This enables organisms to endure severe or persistent environmental stress by altering physiological responses until environmental conditions improve [57, 58] and may facilitate long-term adaptation [46]. Such an adaptive response may also have evolved to allow the females to survive periods of starvation during reproduction and would explain why female lifespan was more robust to the effects of a high carbohydrate evolutionary diet.

A reason for not detecting lifespan extension plasticity on high sugar diets in males might be that these diets represent stressful environments where the optimal lifespan phenotype is constrained, leaving females’ lifespan more receptive to the effects of diets that are optimal for reproduction and lifespan [12, 59, 60]. Indeed, sites of fat deposition in females are different to those of males and have been proposed to reflect adaptations promoting female reproductive success [61]. However, we cannot exclude the possibility that the diet-induced changes might provide other adaptive benefits that trade-off with lifespan and could not be measured easily under laboratory settings, e.g., greater reproductive investment associated with ranging for food that might allow for offspring to survive in poor environments and find more favourable conditions [62].

Although lifespan extension under DR is frequently attributed to energy reallocation for somatic maintenance instead of reproduction [63], the trade-offs sometimes observed between reproduction and lifespan are now thought to relate to hormonal signalling [64] or programmatic mechanisms [65] rather than a shift in reallocation of resources to somatic maintenance from reproduction. Experimental evolution studies have tested this resource reallocation prediction in male and female experimental evolution lines of *D. melanogaster* [23, 24] and shown mixed and sex-specific results. Females evolved shorter lifespan but increased fecundity on DR [24], while males displayed increased fitness at no cost to lifespan on DR [23]. These findings show that lifespan can evolve in response to diet, but the results contradict the typical hypothesis that resource reallocation is driving lifespan extension as predicted under the disposable soma theory of ageing. While examinations of fitness and lifespan trade-offs were beyond the scope of the present study, lifespan extensions have been shown to occur even in sterile and non-reproducing animals in the absence of trade-offs between reproduction and somatic maintenance [66]. While genotypic (female/male) and environmental (sugar rich/sugar poor) specific constraints might also restrict the evolution of phenotypes that are similar in both sexes [23] based on our experimental data, it is reasonable to assume that the effects observed in male and female *D. serrata* under experimental evolution represent similar diet-dependent patterns that have evolved but are different from the plastic lifespan patterns observed in the same populations.

## 5. Conclusion

We previously showed that lifespan variation in *D. serrata* arises due to condition-dependent environmental modulations of underlying genetic variation [67]. Dietary discordance between organisms’ evolutionary and developmental history is increasingly recognised to play a critical role in shaping lifespan and survival [13, 68, 69]. While this study demonstrates that diets with varying levels of carbohydrates can have evolved and plastic effects on lifespan for *D. serrata* on the ancestral dietary environment, there was a pronounced misalignment between the plastic and evolved responses. The match/mismatch implies that plastic responses to high carbohydrates are adaptive but plastic responses to low carbohydrates are not. Here we were only able to test the dietary treatments in a single environment to isolate the evolved response, but whether this holds true across a broader age of evolutionary and developmental diet combinations merits further investigation. This would allow us to test the evolved responses across all the different diets to see if/how plasticity evolves as a correlated response to selection imposed by the different dietary environments. Such match/mismatch approaches could improve the understanding of evolved and plastic responses in natural populations and subsequent consequences to health and lifespan. This approach also helps translate our understanding regarding the impact of contemporary diets into interventions aimed at optimising or ameliorating population health and lifespan [70, 71]. Ultimately, understanding the basis of this variation for lifespan and the different responses to dietary treatments across a broader age of evolutionary and developmental diets may help address the causes and consequences of rapidly changing dietary histories of human populations.

## References

1. Ocampo M., Pincheira-Donoso D., Sayol F., Rios R.S. 2022 Evolutionary transitions in diet influence the exceptional diversification of a lizard adaptive radiation. BMC Ecology and Evolution 22(1), 74. (doi:10.1186/s12862-022-02028-3).

2. Jang T., Lee K.P. 2018 Comparing the impacts of macronutrients on life-history traits in larval and adult *Drosophila melanogaster*: the use of nutritional geometry and chemically defined diets. The Journal of experimental biology 221(Pt 21). (doi:10.1242/jeb.181115).

3. Lee D., Hwang W., Artan M., Jeong D.E., Lee S.J. 2015 Effects of nutritional components on aging. Aging Cell 14(1), 8–16. (doi:10.1111/acel.12277).

4. Lee D., Son H.G., Jung Y., Lee S.V. 2017 The role of dietary carbohydrates in organismal aging. Cell Mol Life Sci 74(10), 1793–1803. (doi:10.1007/s00018-016-2432-6).

5. Lee K.P. 2015 Dietary protein:carbohydrate balance is a critical modulator of lifespan and reproduction in *Drosophila melanogaster*: a test using a chemically defined diet. J Insect Physiol 75, 12–19. (doi:10.1016/j.jinsphys.2015.02.007).

6. Min K.W., Jang T., Lee K.P. 2021 Thermal and nutritional environments during development exert different effects on adult reproductive success in *Drosophila melanogaster*. Ecol Evol 11(1), 443–457. (doi:10.1002/ece3.7064).

7. Zajitschek F., Jin T., Colchero F., Maklakov A.A. 2014 Aging differently: diet-and sex-dependent late-life mortality patterns in *Drosophila melanogaster*. J Gerontol A Biol Sci Med Sci 69(6), 666–674. (doi:10.1093/gerona/glt158).

8. Archer C.R., Basellini U., Hunt J., Simpson S.J., Lee K.P., Baudisch A. 2018 Diet has independent effects on the pace and shape of aging in *Drosophila melanogaster*. Biogerontology 19(1), 1–12. (doi:10.1007/s10522-017-9729-1).

9. Barson N.J., Aykanat T., Hindar K., Baranski M., Bolstad G.H., Fiske P., Jacq C., Jensen A.J., Johnston S.E., Karlsson S., et al. 2015 Sex-dependent dominance at a single locus maintains variation in age at maturity in salmon. Nature 528(7582), 405–408. (doi:10.1038/nature16062).

10. Brengdahl M., Kimber C.M., Maguire-Baxter J., Malacrino A., Friberg U. 2018 Genetic Quality Affects the Rate of Male and Female Reproductive Aging Differently in *Drosophila melanogaster*. Am Nat 192(6), 761–772. (doi:10.1086/700117).

11. Camus M.F., Fowler K., Piper M.W.D., Reuter M. 2017 Sex and genotype effects on nutrient-dependent fitness landscapes in *Drosophila melanogaster*. Proc Biol Sci 284(1869). (doi:10.1098/rspb.2017.2237).

12. Camus M.F., Piper M.D., Reuter M. 2019 Sex-specific transcriptomic responses to changes in the nutritional environment. Elife 8. (doi:10.7554/eLife.47262).

13. Duxbury E.M.L., Chapman T. 2019 Sex-Specific Responses of Lifespan and Fitness to Variation in Developmental versus Adult Diets in D. melanogaster. J Gerontol A Biol Sci Med Sci. (doi:10.1093/gerona/glz175).

14. Hosking C.J., Raubenheimer D., Charleston M.A., Simpson S.J., Senior A.M. 2019 Macronutrient intakes and the lifespan-fecundity trade-off: a geometric framework agent-based model. J R Soc Interface 16(151), 20180733. (doi:10.1098/rsif.2018.0733).

15. Jensen K., McClure C., Priest N.K., Hunt J. 2015 Sex-specific effects of protein and carbohydrate intake on reproduction but not lifespan in *Drosophila melanogaster*. Aging Cell 14(4), 605–615. (doi:10.1111/acel.12333).

16. Klepsatel P., Knoblochova D., Girish T.N., Dircksen H., Galikova M. 2020 The influence of developmental diet on reproduction and metabolism in *Drosophila*. BMC Evol Biol 20(1), 93. (doi:10.1186/s12862-020-01663-y).

17. Lihoreau M., Poissonnier L.A., Isabel G., Dussutour A. 2016 *Drosophila* females trade off good nutrition with high-quality oviposition sites when choosing foods. The Journal of experimental biology 219(Pt 16), 2514–2524. (doi:10.1242/jeb.142257).

18. Dobson A.J., Ezcurra M., Flanagan C.E., Summerfield A.C., Piper M.D.W., Gems D., Alic N. 2017 Nutritional Programming of Lifespan by FOXO Inhibition on Sugar-Rich Diets. Cell Rep 18(2), 299–306. (doi:10.1016/j.celrep.2016.12.029).

19. Emborski C., Mikheyev A.S. 2019 Ancestral diet transgenerationally influences offspring in a parent-of-origin and sex-specific manner. Philos Trans R Soc Lond B Biol Sci 374(1768), 20180181. (doi:10.1098/rstb.2018.0181).

20. Furse S., Watkins A.J., Hojat N., Smith J., Williams H.E.L., Chiarugi D., Koulman A. 2021 Lipid Traffic Analysis reveals the impact of high paternal carbohydrate intake on offsprings’ lipid metabolism. Commun Biol 4(1), 163. (doi:10.1038/s42003-021-01686-1).

21. Ivimey-Cook E.R., Sales K., Carlsson H., Immler S., Chapman T., Maklakov A.A. 2021 Transgenerational fitness effects of lifespan extension by dietary restriction in Caenorhabditis elegans. Proc Biol Sci 288(1950), 20210701. (doi:10.1098/rspb.2021.0701).

22. Öst A., Lempradl A., Casas E., Weigert M., Tiko T., Deniz M., Pantano L., Boenisch U., Itskov P.M., Stoeckius M., et al. 2014 Paternal diet defines offspring chromatin state and intergenerational obesity. Cell 159(6), 1352–1364. (doi:10.1016/j.cell.2014.11.005).

23. Zajitschek F., Georgolopoulos G., Vourlou A., Ericsson M., Zajitschek S.R.K., Friberg U., Maklakov A. A. 2019 Evolution Under Dietary Restriction Decouples Survival From Fecundity in *Drosophila melanogaster* Females. J Gerontol A Biol Sci Med Sci 74(10), 1542–1548. (doi:10.1093/gerona/gly070).

24. Zajitschek F., Zajitschek S.R., Canton C., Georgolopoulos G., Friberg U., Maklakov A.A. 2016 Evolution under dietary restriction increases male reproductive performance without survival cost. Proc Biol Sci 283(1825), 20152726. (doi:10.1098/rspb.2015.2726).

25. Vinton A.C., Gascoigne S.J.L., Sepil I., Salguero-Gomez R. 2022 Plasticity’s role in adaptive evolution depends on environmental change components. Trends Ecol Evol. (doi:10.1016/j.tree.2022.08.008).

26. Ghalambor C.K., McKay J.K., Carroll S.P., Reznick D.N. 2007 Adaptive versus non-adaptive phenotypic plasticity and the potential for contemporary adaptation in new environments. Functional Ecology 21(3), 394–407. (doi:https://doi.org/10.1111/j.1365-2435.2007.01283.x).

27. Chevin L.-M., Lande R., Mace G.M. 2010 Adaptation, Plasticity, and Extinction in a Changing Environment: Towards a Predictive Theory. PLOS Biology 8(4), e1000357. (doi:10.1371/journal.pbio.1000357).

28. Pigliucci M., Murren C.J., Schlichting C.D. 2006 Phenotypic plasticity and evolution by genetic assimilation. Journal of Experimental Biology 209(12), 2362–2367.

29. Grether G.F. 2005 Environmental Change, Phenotypic Plasticity, and Genetic Compensation. The American Naturalist 166(4), E115–E123. (doi:10.1086/432023).

30. Schaum C.E., Collins S. 2014 Plasticity predicts evolution in a marine alga. Proc Biol Sci 281(1793). (doi:10.1098/rspb.2014.1486).

31. Lucas A. 1998 Programming by Early Nutrition: An Experimental Approach. The Journal of Nutrition 128(2), 401S-406S. (doi:10.1093/jn/128.2.401S).

32. Baugh L.R., Day T. 2020 Nongenetic inheritance and multigenerational plasticity in the nematode C. elegans. Elife 9. (doi:10.7554/eLife.58498).

33. Mair W., Goymer P., Pletcher S.D., Partridge L. 2003 Demography of dietary restriction and death in *Drosophila*. Science 301(5640), 1731–1733. (doi:10.1126/science.1086016).

34. Simons M.J., Koch W., Verhulst S. 2013 Dietary restriction of rodents decreases aging rate without affecting initial mortality rate -- a meta-analysis. Aging Cell 12(3), 410–414. (doi:10.1111/acel.12061).

35. McCracken A.W., Buckle E., Simons M.J.P. 2020 The relationship between longevity and diet is genotype dependent and sensitive to desiccation in *Drosophila melanogaster*. The Journal of experimental biology 223(Pt 23). (doi:10.1242/jeb.230185).

36. Reddiex A.J., Allen S.L., Chenoweth S.F. 2018 A Genomic Reference Panel for *Drosophila serrata*. G3 (Bethesda) 8(4), 1335–1346. (doi:10.1534/g3.117.300487).

37. Littell R.C., Milliken G.A., Stroup W.W., Wolfinger R.D., Oliver S. 2006 SAS for mixed models, SAS publishing.

38. Stanley P.D., Ng’oma E., O’Day S., King E.G. 2017 Genetic Dissection of Nutrition-Induced Plasticity in Insulin/Insulin-Like Growth Factor Signaling and Median Life Span in a *Drosophila* Multiparent Population. Genetics 206(2), 587–602. (doi:10.1534/genetics.116.197780).

39. Cordain L., Eaton S.B., Sebastian A., Mann N., Lindeberg S., Watkins B.A., O’Keefe J.H., Brand-Miller J. 2005 Origins and evolution of the Western diet: health implications for the 21st century. Am J Clin Nutr 81(2), 341–354. (doi:10.1093/ajcn.81.2.341).

40. Luca F., Perry G.H., Di Rienzo A. 2010 Evolutionary adaptations to dietary changes. Annu Rev Nutr 30, 291–314. (doi:10.1146/annurev-nutr-080508-141048).

41. Skorupa D.A., Dervisefendic A., Zwiener J., Pletcher S.D. 2008 Dietary composition specifies consumption, obesity, and lifespan in *Drosophila melanogaster*. Aging Cell 7(4), 478–490. (doi:10.1111/j.1474-9726.2008.00400.x).

42. Chandegra B., Tang J.L.Y., Chi H., Alic N. 2017 Sexually dimorphic effects of dietary sugar on lifespan, feeding and starvation resistance in *Drosophila*. Aging 9(12), 2521–2528. (doi:10.18632/aging.101335).

43. Huang Y., Agrawal A.F. 2016 Experimental Evolution of Gene Expression and Plasticity in Alternative Selective Regimes. PLoS Genet 12(9), e1006336. (doi:10.1371/journal.pgen.1006336).

44. Agrawal A.F., Whitlock M.C. 2010 Environmental duress and epistasis: how does stress affect the strength of selection on new mutations? Trends in ecology & evolution 25(8), 450–458.

45. Hoffmann A.A., Hercus M.J. 2000 Environmental stress as an evolutionary force. Bioscience 50(3), 217–226.

46. Lande R. 2009 Adaptation to an extraordinary environment by evolution of phenotypic plasticity and genetic assimilation. Journal of evolutionary biology 22(7), 1435–1446.

47. Rohner P.T., Teder T., Esperk T., Lüpold S., Blanckenhorn W.U. 2018 The evolution of male-biased sexual size dimorphism is associated with increased body size plasticity in males. Functional ecology 32(2), 581–591.

48. Ormerod K.G., LePine O.K., Abbineni P.S., Bridgeman J.M., Coorssen J.R., Mercier A.J., Tattersall G.J. 2017 *Drosophila* development, physiology, behavior, and lifespan are influenced by altered dietary composition. Fly (Austin) 11(3), 153–170. (doi:10.1080/19336934.2017.1304331).

49. Wu Q., Yu G., Cheng X., Gao Y., Fan X., Yang D., Xie M., Wang T., Piper M.D.W., Yang M. 2020 Sexual dimorphism in the nutritional requirement for adult lifespan in *Drosophila melanogaster*. Aging Cell 19(3), e13120. (doi:10.1111/acel.13120).

50. Chen H.Y., Maklakov A.A. 2012 Longer life span evolves under high rates of condition-dependent mortality. Curr Biol 22(22), 2140–2143. (doi:10.1016/j.cub.2012.09.021).

51. Rattan S.I., Fernandes R.A., Demirovic D., Dymek B., Lima C.F. 2009 Heat stress and hormetin-induced hormesis in human cells: effects on aging, wound healing, angiogenesis, and differentiation. Dose Response 7(1), 90–103. (doi:10.2203/dose-response.08-014.Rattan).

52. Sinclair D.A. 2005 Toward a unified theory of caloric restriction and longevity regulation. Mech Ageing Dev 126(9), 987–1002. (doi:10.1016/j.mad.2005.03.019).

53. Giugliano D., Maiorino M.I., Bellastella G., Esposito K. 2018 More sugar? No, thank you! The elusive nature of low carbohydrate diets. Endocrine 61(3), 383–387. (doi:10.1007/s12020-018-1580-x).

54. Bruce K.D., Hoxha S., Carvalho G.B., Yamada R., Wang H.D., Karayan P., He S., Brummel T., Kapahi P., Ja W.W. 2013 High carbohydrate-low protein consumption maximizes *Drosophila* lifespan. Experimental gerontology 48(10), 1129–1135. (doi:10.1016/j.exger.2013.02.003).

55. Shokhirev M.N., Johnson A.A. 2014 Effects of extrinsic mortality on the evolution of aging: a stochastic modeling approach. PloS one 9(1), e86602. (doi:10.1371/journal.pone.0086602).

56. Lafuente E., Beldade P. 2019 Genomics of developmental plasticity in animals. Frontiers in genetics 10, 720.

57. Braendle C., Félix M.-A. 2008 Plasticity and errors of a robust developmental system in different environments. Developmental cell 15(5), 714–724.

58. Sommer R.J. 2020 Phenotypic plasticity: from theory and genetics to current and future challenges. Genetics 215(1), 1–13.

59. Kim K.E., Jang T., Lee K.P. 2020 Combined effects of temperature and macronutrient balance on life-history traits in *Drosophila melanogaster*: implications for life-history trade-offs and fundamental niche. Oecologia 193(2), 299–309. (doi:10.1007/s00442-020-04666-0).

60. Rapkin J., Jensen K., Archer C.R., House C.M., Sakaluk S.K., Castillo E.D., Hunt J. 2018 The Geometry of Nutrient Space-Based Life-History Trade-Offs: Sex-Specific Effects of Macronutrient Intake on the Trade-Off between Encapsulation Ability and Reproductive Effort in Decorated Crickets. Am Nat 191(4), 452–474. (doi:10.1086/696147).

61. Power M.L., Schulkin J. 2008 Sex differences in fat storage, fat metabolism, and the health risks from obesity: possible evolutionary origins. Br J Nutr 99(5), 931–940. (doi:10.1017/s0007114507853347).

62. Pontzer H., Kamilar J.M. 2009 Great ranging associated with greater reproductive investment in mammals. Proceedings of the National Academy of Sciences 106(1), 192–196. (doi:10.1073/pnas.0806105106).

63. Kirkwood T.B. 1977 Evolution of ageing. Nature 270(5635), 301–304.

64. Adler M.I., Cassidy E.J., Fricke C., Bonduriansky R. 2013 The lifespan-reproduction trade-off under dietary restriction is sex-specific and context-dependent. Experimental gerontology 48(6), 539–548. (doi:10.1016/j.exger.2013.03.007).

65. Gems D. 2022 The hyperfunction theory: An emerging paradigm for the biology of aging. Ageing Res Rev 74, 101557. (doi:10.1016/j.arr.2021.101557).

66. Ghazi A., Henis-Korenblit S., Kenyon C. 2009 A transcription elongation factor that links signals from the reproductive system to lifespan extension in Caenorhabditis elegans. PLoS Genet 5(9), e1000639. (doi:10.1371/journal.pgen.1000639).

67. Narayan V.P., Wilson A.J., Chenoweth S.F. 2022 Genetic and social contributions to sex differences in lifespan in *Drosophila serrata*. J Evol Biol. (doi:10.1111/jeb.13992).

68. Feige-Diller J., Palme R., Kaiser S., Sachser N., Richter S.H. 2021 The impact of varying food availability on health and welfare in mice: Testing the Match-Mismatch hypothesis. Physiol Behav 228, 113193. (doi:10.1016/j.physbeh.2020.113193).

69. Raubenheimer D., Simpson S.J., Tait A.H. 2012 Match and mismatch: conservation physiology, nutritional ecology and the timescales of biological adaptation. Philos Trans R Soc Lond B Biol Sci 367(1596), 1628–1646. (doi:10.1098/rstb.2012.0007).

70. Monaghan P. 2008 Early growth conditions, phenotypic development and environmental change. Philos Trans R Soc Lond B Biol Sci 363(1497), 1635–1645. (doi:10.1098/rstb.2007.0011).

71. Tilman D., Clark M. 2014 Global diets link environmental sustainability and human health. Nature 515(7528), 518–522. (doi:10.1038/nature13959).

