## Supplementary Tables 1-7 and Supplemetary Figures 1-2. for "Misaligned plastic and evolutionary responses of lifespan to novel carbohydrate diets"

**Table S1 Diet composition**

| S:Y ratio | 1:0.5 | 1:1.0 | 1:1.5 | 1:2.0 | 1:2.5 |
|---|---|---|---|---|---|
| Agar (g/L) | 18.0 | 18.0 | 18.0 | 18.0 | 18.0 |
| Sucrose (g/L) | 18.5 | 37.0 | 54.0 | 74.0 | 92.5 |
| Brewer's yeast (g/L) | 37.0 | 37.0 | 37.0 | 37.0 | 37.0 |
| Nipagin solution* (ml/L) | 12.0 | 12.0 | 12.0 | 12.0 | 12.0 |
| Propionic acid (ml/L) | 6.0 | 6.0 | 6.0 | 6.0 | 6.0 |

For each diet: 1L of distilled water was added.

**Table S2 Summary statistics for lifespan on the 1.5 SY Developmental diet**

| | Females | | Males | |
|---|---|---|---|---|
| **Evolutionary Diet** | **Mean ± SE** | **Sample size** | **Mean ± SE** | **Sample size** |
| 0.5 SY | 72.7 ± 1.44 | 386.0 | 50.4 ± 1.59 | 391.0 |
| 1.0 SY | 72.7 ± 1.43 | 385.0 | 50.4 ± 1.59 | 391.0 |
| 1.5 SY | 67.4 ± 1.32 | 385.0 | 48.0 ± 1.85 | 389.0 |
| 2.0 SY | 66.9 ± 1.64 | 391.0 | 40.2 ± 1.62 | 390.0 |
| 2.5 SY | 63.2 ± 1.99 | 388.0 | 40.8 ± 2.03 | 393.0 |

**Table S3 Summary statistics for lifespan of the 1.5 SY Evolutionary diet**

| | Females | | Males | |
|---|---|---|---|---|
| **Developmental Diet** | **Mean ± SE** | **Sample size** | **Mean ± SE** | **Sample size** |
| 0.5 SY | 62.2 ± 0.99 | 386.0 | 44.0 ± 1.59 | 394.0 |
| 1.0 SY | 65.2 ± 0.80 | 392.0 | 45.2 ± 2.80 | 399.0 |
| 1.5 SY | 67.4 ± 1.33 | 385.0 | 48.0 ± 1.86 | 389.0 |
| 2.0 SY | 64.7 ± 1.45 | 392.0 | 42.9 ± 1.44 | 385.0 |
| 2.5 SY | 56.8 ± 1.25 | 396.0 | 42.0 ± 1.65 | 380.0 |

**Table S4 post-hoc tests for effects of Evolutionary diet on female lifespan.** Values are least square estimates of the fixed effect 'Developmental diet' in the final model, containing 'Evolutionary diet' as fixed effects, without their interaction term, i.e., controlling for Developmental diet.

| Sex (Assay Diet) | Evolution Diet Comparisons | | Estimate | Standard Error | DF | t Value | Pr > \|t\| | Adj P |
|---|---|---|---|---|---|---|---|---|
| Females (1.5 SY) | 0.5 SY | 1.5 SY | 5.11 | 2.49 | 38.3 | 2.05 | **0.05** | 0.24 |
| | 1.0 SY | 1.5 SY | 5.15 | 2.50 | 38.4 | 2.06 | **0.05** | 0.24 |
| | 1.5 SY | 2.0 SY | 0.62 | 2.49 | 38.1 | 0.25 | 0.80 | 1.00 |
| | 1.5 SY | 2.5 SY | 4.41 | 2.49 | 38.1 | 1.77 | 0.09 | 0.39 |

**Table S5 post-hoc tests for effects of Evolutionary diet on male lifespan.** Values are least square estimates of the fixed effect 'Developmental diet' in the final model, containing 'Evolutionary diet' as fixed effects, without their interaction term, i.e., controlling for Developmental diet

| Sex (Assay Diet) | Evolution Diet Comparisons | | Estimate | Standard Error | DF | t Value | Pr > \|t\| | Adj P |
|---|---|---|---|---|---|---|---|---|
| Males (1.5 SY) | 0.5 SY | 1.5 SY | 2.21 | 2.49 | 37.9 | 0.89 | 0.38 | 0.90 |
| | 1.0 SY | 1.5 SY | 2.21 | 2.49 | 37.9 | 0.89 | 0.38 | 0.90 |
| | 1.5 SY | 2.0 SY | 7.94 | 2.49 | 38.0 | 3.19 | **0.003** | 0.01 |
| | 1.5 SY | 2.5 SY | 7.48 | 2.49 | 37.9 | 3.01 | **0.005** | 0.02 |

**Table S6 post-hoc tests for effects of Developmental diet on female lifespan.** Values are least square estimates of the fixed effect 'Evolutionary diet' in the final model, containing 'Developmental diet' as fixed effects, without their interaction term, i.e., controlling for Evolutionary diet.

| **Sex (Developmental Diet)** | **Evolutionary Diet Comparisons** | | **Estimate** | **Standard Error** | **DF** | **t Value** | **Pr > \|t\|** | **Adj P** |
|---|---|---|---|---|---|---|---|---|
| Females (1.5 SY) | 0.5 SY | 1.5 SY | -5.33 | 2.20 | 32.0 | -2.42 | **0.02** | 0.11 |
| | 1.0 SY | 1.5 SY | -2.32 | 2.19 | 31.7 | -1.06 | 0.30 | 0.83 |
| | 1.5 SY | 2.0 SY | 2.76 | 2.19 | 31.7 | 1.26 | 0.22 | 0.72 |
| | 1.5 SY | 2.5 SY | 10.7 | 2.19 | 31.6 | 4.90 | **<.0001** | <.0001 |

**Table S7 post-hoc tests for effects of Developmental diet on male lifespan.** Values are least square estimates of the fixed effect 'Evolutionary diet' in the final model, containing 'Developmental diet' as fixed effects, without their interaction term, i.e., controlling for Evolutionary diet.

| **Sex (Developmental Diet)** | **Evolutionary Diet Comparisons** | | **Estimate** | **Standard Error** | **DF** | **t Value** | **Pr > \|t\|** | **Adj P** |
|---|---|---|---|---|---|---|---|---|
| Males (1.5 SY) | 0.5 SY | 1.5 SY | -4.17 | 2.19 | 31.6 | -1.90 | 0.07 | 0.32 |
| | 1.0 SY | 1.5 SY | -3.01 | 2.19 | 31.4 | -1.38 | 0.18 | 0.64 |
| | 1.5 SY | 2.0 SY | 5.30 | 2.19 | 31.9 | 2.41 | **0.02** | 0.11 |
| | 1.5 SY | 2.5 SY | 6.26 | 2.20 | 32.0 | 2.85 | **0.01** | 0.04 |

A )

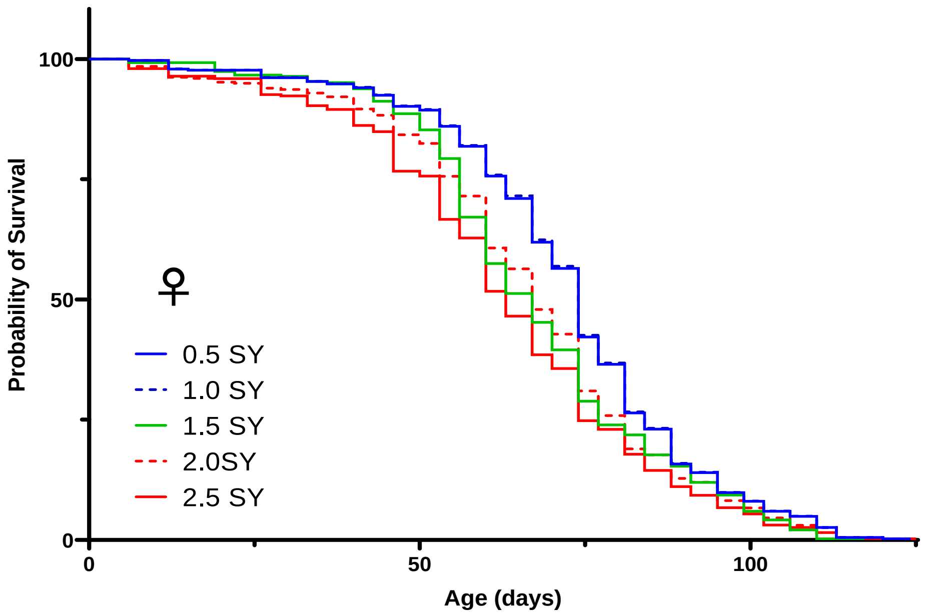


B )

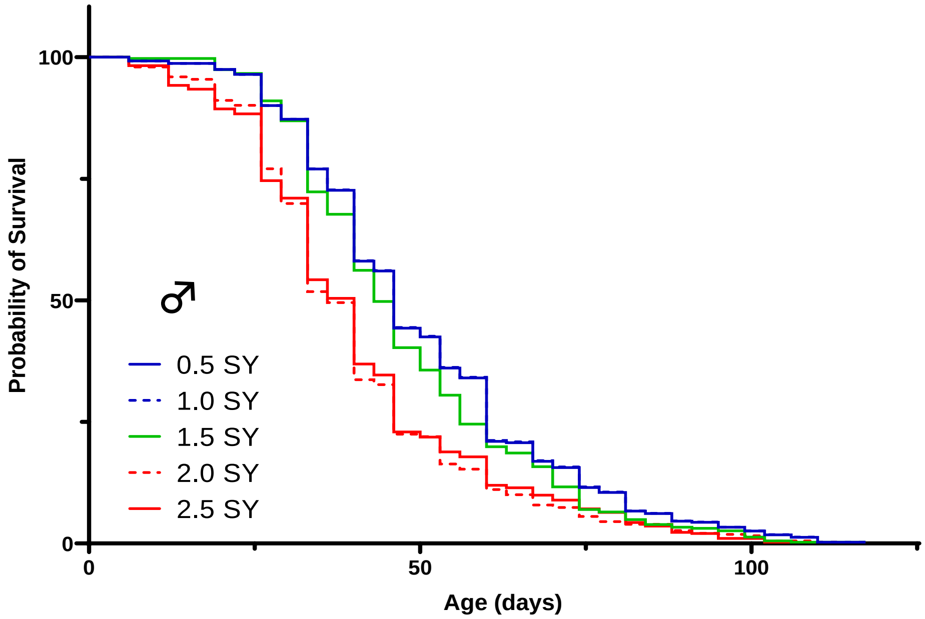


**Figure S1 Survivorship of *D. serrata* offspring transferred back to the ancestral diet of 1.5 SY.** Each panel shows Kaplan–Meier survival curves for A) female and B) male flies. Survival was plotted as adult lifespan in days with where separate curves depict survivorship of evolutionary diet populations 0.5 SY, 1.0 SY, 1.5 SY, 2.0 SY, and 2.5 SY, tested on 1.5 SY developmental diet.

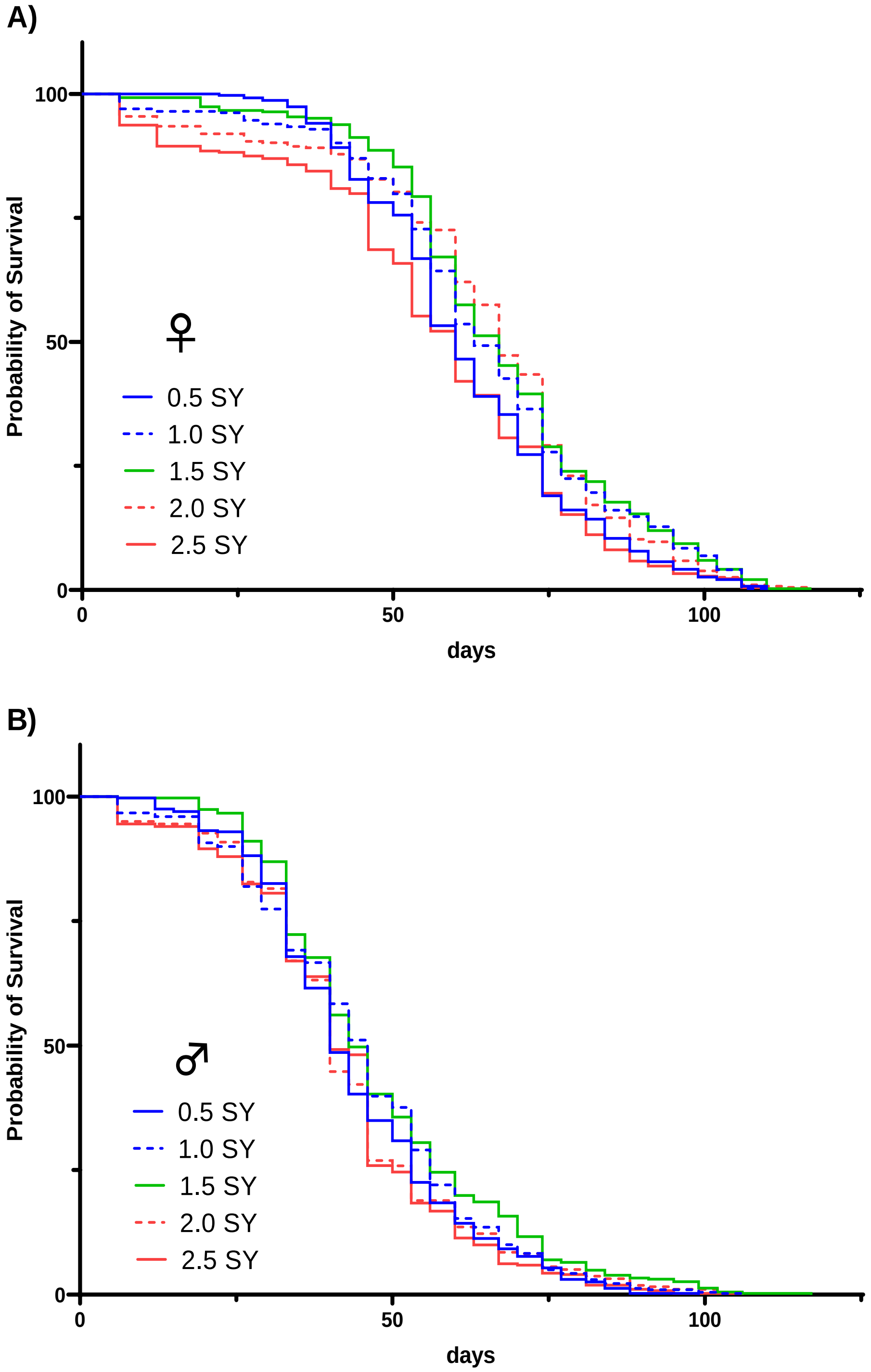


**Figure S2 Survivorship of *D. serrata* offspring transferred from the ancestral diet to every other developmental diet.** Each panel shows Kaplan–Meier survival curves for A) female and B) male flies. Survival was plotted as adult lifespan in days with where separate curves depict survivorship of developmental diet populations 0.5 SY, 1.0 SY, 1.5 SY, 2.0 SY, and 2.5 SY, tested on 1.5 SY ancestral diet.
